# Linking soil biology and chemistry using bacterial isolate exometabolite profiles

**DOI:** 10.1101/109330

**Authors:** Tami L. Swenson, Ulas Karaoz, Joel M. Swenson, Benjamin P. Bowen, Trent Northen

## Abstract

Sequencing provides a window into microbial community structure and metabolic potential; however, linking these data to exogenous metabolites that microorganisms process and produce (the exometabolome) remains challenging. Previously, we observed strong exometabolite niche partitioning among bacterial isolates from biological soil crust (biocrust). Here we examine native biocrust to determine if these patterns are reproduced in the environment. Overall, most soil metabolites displayed the expected relationship (positive or negative correlation) with four dominant bacteria following a wetting event and across biocrust developmental stages. For metabolites that were previously found to be consumed by an isolate, 78% were negatively correlated with the abundance of *in situ* isolate phylotypes whereas for released metabolites, 73% were positively correlated. Our results demonstrate that metabolite profiling, sequencing and exometabolomics can be successfully integrated to functionally link metagenomes and microbial community structure with environmental chemistry.

---

In soils, which harbor the largest terrestrial pool of organic carbon^1^, organic matter is largely processed by complex microbial communities. The impact of climate change on these communities and their activities is uncertain^2^. Given the importance of these systems, vast amounts of sequencing data have been and continue to be collected. While metagenomic sequencing provides important insights into community structure and metabolic potential, if unconstrained, such data are often open to multiple interpretations. New approaches are needed to help link the now readily available sequencing data to *in situ* metabolism in order to better understand the dynamic reciprocity between carbon cycling and microbial community structure.

Soil organic matter (SOM) content and moisture have long been recognized as important factors controlling soil microbial community structure and carbon cycling^3,4^. For example, microbial community diversity and richness are positively correlated with soil organics across diverse ecosystems including polar soils^5^, agricultural soils^6^ and arid soils^7^. Similarly, soil wetting events are well-known to dramatically alter community structure^8^ including establishing cascades of microbial abundances^9^. Arid lands account for over 40% of Earth’s terrestrial surface^10^ and are especially sensitive to SOM and moisture content. It is predicted that the aridity of drylands will increase, reducing SOM and microbial community diversity, and that this will impact ecosystem productivity^7,11^. This strong coupling between soil moisture, SOM and community structure is especially important in the arid land topsoil microbial communities known as biological soil crusts (biocrusts), which cover a large fraction of arid regions and are critical in nutrient cycling^12^. Biocrusts exist in a dormant desiccated state and only become metabolically active during infrequent rainfall events^13^ and like other soils, organic matter plays a vital role in retaining moisture and increasing microbial diversity^14^.

The mechanisms linking SOM composition and microbial community structure are poorly understood. It is now thought that the organic matter that is cycled by soil microbes is a complex mixture of microbial metabolites^15,16^ that can be characterized in detail using soil metabolomics^17,18^. The composition of these exometabolites has a large impact on community structure, and in turn, these microbes impact the metabolite pool. For example, in some cases, resource competition can reduce diversity through competitive exclusion, whereas cross-feeding can increase diversity. On the other hand, rich sources of SOM may promote diversity through niche divergence^19^ and exometabolite niche partitioning^20^.

Exometabolomics enables direct examination of how microbes transform the small molecule metabolites within their environment, providing new insights into resource competition and cross-feeding^21^. For this approach, microbes (typically isolates) are cultured in an environmentally-relevant mixture of metabolites and then spent media is profiled to determine the uptake and release of metabolites. Recently, exometabolomics was used to study resource partitioning among sympatric biocrust isolates using complex media^20^. This revealed a high degree of substrate specialization where 13-26% of the detected metabolites were consumed by individual isolates. As organisms from diverse taxa continue to be cultivated and examined, this approach holds substantial potential to provide valuable phenotypic information that can link community structure to SOM composition.

Here we explored the dynamics and relationships between biocrust microbes and metabolites. We then determined the extent to which isolate exometabolite patterns are conserved *in situ* (within the intact soil community). Microbe-metabolite changes were driven by wetting dry biocrusts obtained along an ecological successional gradient (Figure 1A). Four successional stages of biocrust were used, ranging from early/ young (labeled as ‘level A’) to late/ mature (labeled as ‘level D’) (Supplementary Figure 1). We then compared our current results with previous laboratory-derived knowledge of substrate preferences for four dominant organisms by relating the abundance of these bacteria to soil metabolites measured in the intact biocrust system (Figure 1B). While the comparison of a microbe in isolation and in an environmental system is complex, the general assumption is that as a particular microbe grows and increases in abundance in a community, consumed metabolites will decrease and display a negative-correlation relationship. Conversely, metabolites that are known to be released by a microbe are predicted to concurrently increase and display a positive-correlation relationship with growth (Figure 1B). Liquid chromatography-mass spectrometry (LC/MS) soil metabolomics was used to characterize the dynamic composition of the biocrust soil water and shotgun metagenomic sequencing was used to measure single copy gene markers of the dominant taxa. To the best of our knowledge, this is the first study using isolate exometabolomics to link microbial community structure to soil chemistry.

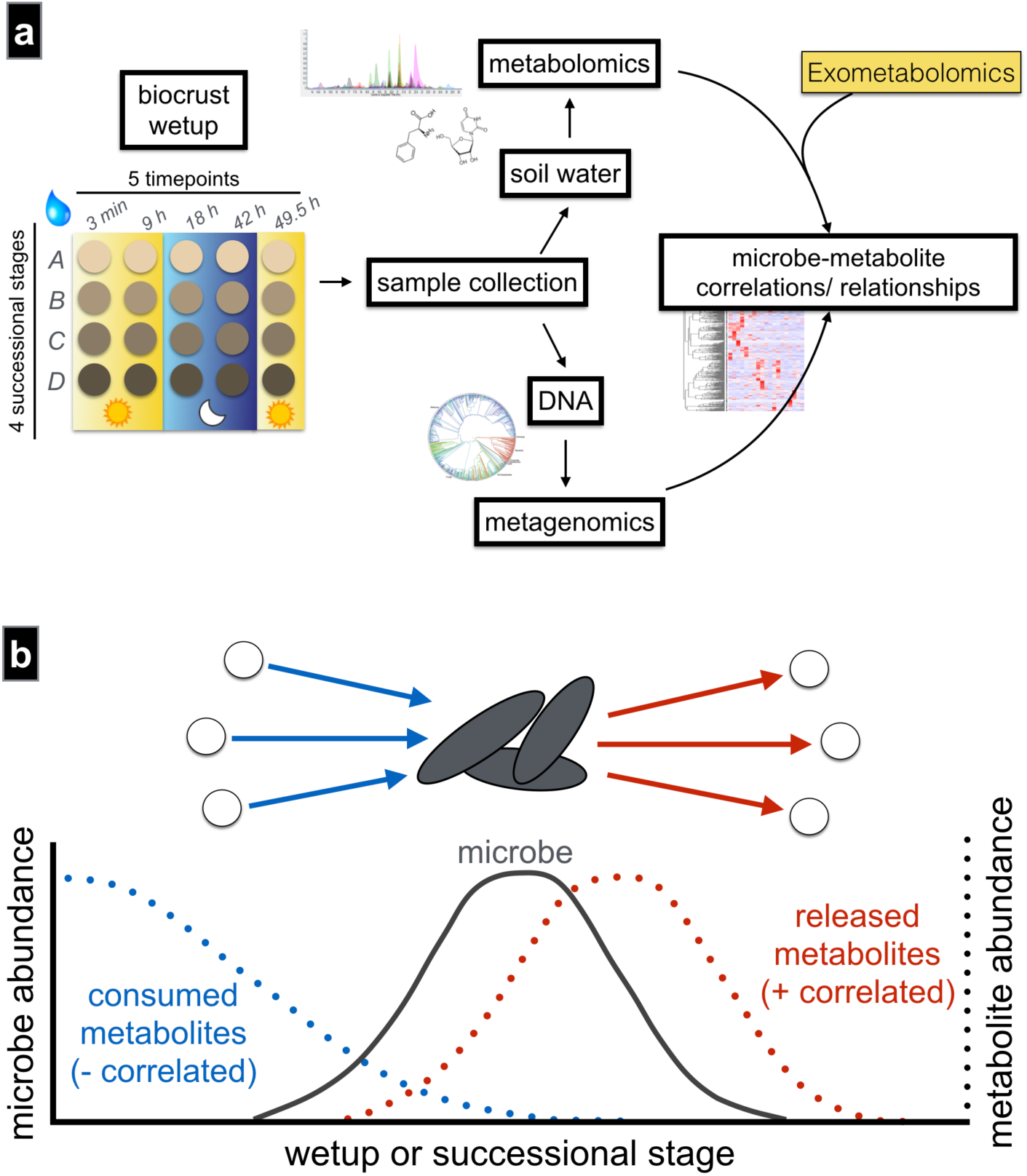
A. Biocrust wetup metabolomics and metagenomics experimental setup and analysis. To study microbe-metabolite relationships *in situ*, biocrusts from four successional stages were wetup and sampled at five time points. Biocrust soil water was removed and analyzed by liquid chromatography/ mass spectrometry and biocrust DNA was extracted for sequencing. Metagenome-estimated genome and metabolite abundances were analyzed through rank correlations to determine microbe-metabolite relationships and compared to the expected relationships based on isolate exometabolomic studies. **B. Exometabolomics-based *in situ* microbe-metabolite relationship prediction.** The hypothesis is that isolate exometabolomics can be used to predict microbe-metabolite patterns *in situ* based on microbial abundance: Across wetting and successional stages, microbes change in abundance and negatively correlate with metabolites that they consume and positively correlate with metabolites that they release (metabolites are indicated by dotted lines).

## RESULTS

### Cycling of metabolites and microbes across wetting and successional stages

The metabolic activity caused by wetting was monitored at various time points ranging from immediate (3 min) to long-term (49.5 h) across the four biocrust successional stages. Biocrust soil water was analyzed by LC/MS, resulting in the identification of 85 metabolites using authentic chemical standards (Figure 2 and Supplementary Table 1). All metabolites displayed cycling by changing at least two-fold (between minimum and maximum peak areas) across both wetting and successional stages (Figure 2). Wetting duration had a stronger impact on metabolite dynamics compared to changes in successional stages (Supplementary Figure 2). Hierarchical clustering of metabolite patterns revealed three distinct clusters (Figure 2). The first cluster (cluster 1, Figure 2) consisted of most (5 out of 7) of the detected fatty acids (palmitate, myristate, stearate, laurate, decanoate), which were most abundant at the first time point (3 min) for all successional stages, and gradually decreased with time. The largest cluster (cluster 2, Figure 2) was enriched with the majority of amino acids and nucleobases, which peaked in abundance at the early to early-mid time points. Within this cluster, the earliest metabolites included polar amino acids (glutamine, glutamate, asparagine, 4-oxoproline, aspartate and lysine) and the nucleobases uridine, guanosine and cytidine. The final cluster (cluster 3 in Figure 2) contained metabolites most abundant at late time points and in more mature biocrust (*e.g.* salicylate, panthothenate, nicotinate, xanthine, creatinine).

**Figure 2.**
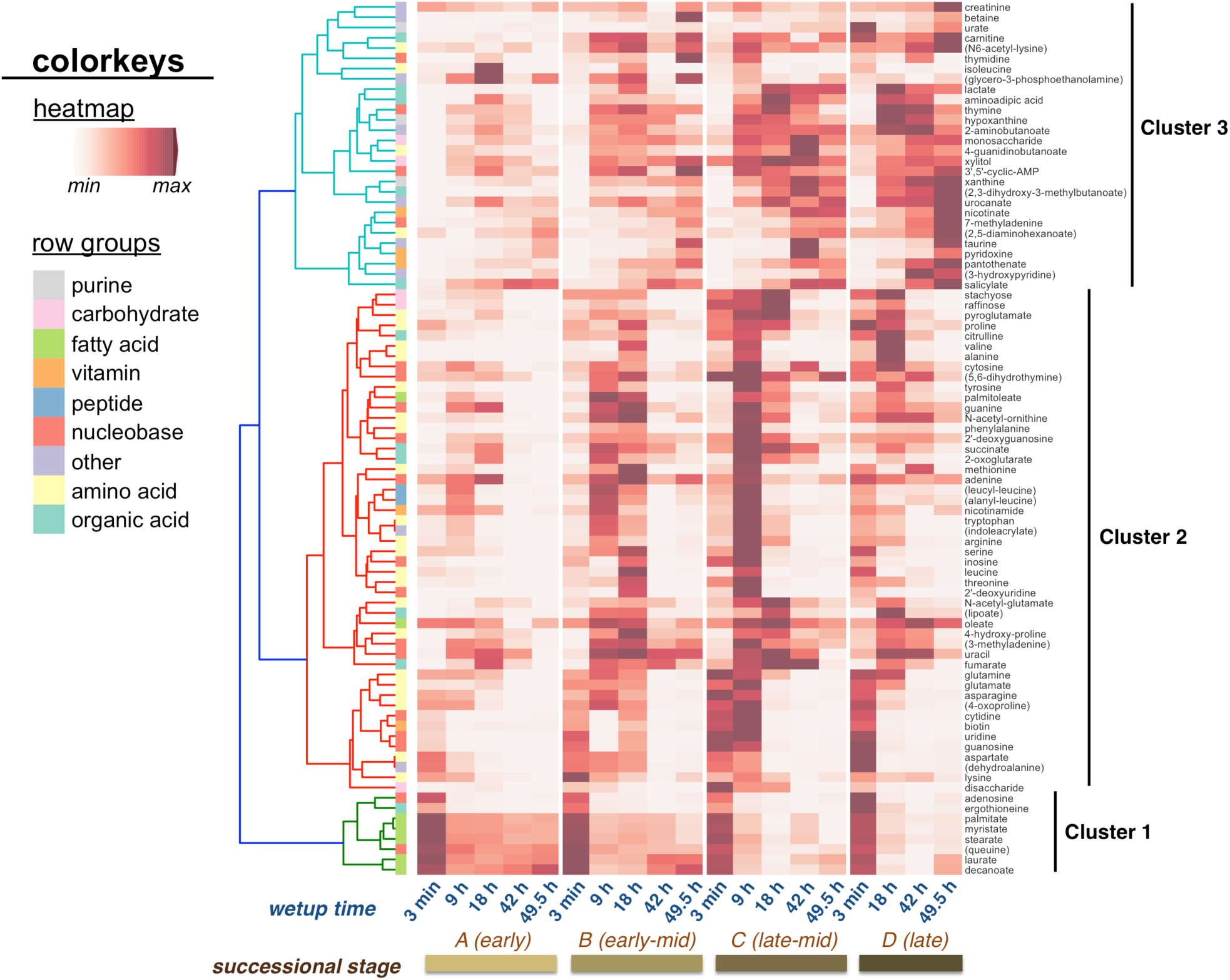
Metabolite patterns detected in biocrust soil water. Metabolite cycling, for a total of 85 metabolites, was observed in biocrust soil water across wetting and successional stages. Unique patterns are indicated by cluster 1 (early metabolites, fatty acids), cluster 2 (early-to mid-time point metabolites) and cluster 3 (late metabolites). Putative metabolites are indicated by parentheses.

**Table 1.**
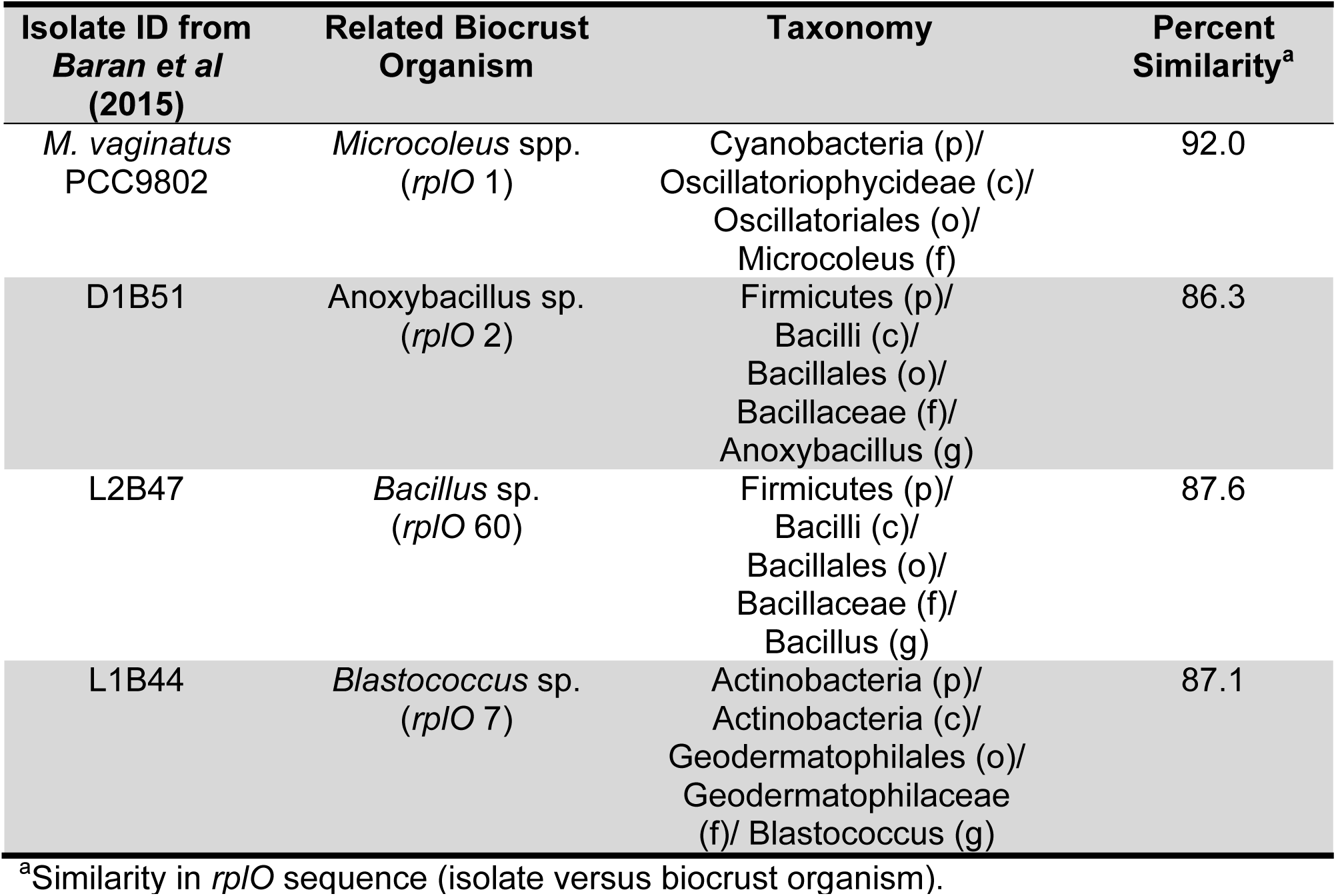
Exometabolite-profiled isolates^20^ and their closest phylotypes in biocrust.

Microbial community structure was determined using shotgun metagenomics with a genome-centric analysis pipeline. The relative abundances of environmental genomes were determined via read-mapping to a universal single-copy phylogenetic marker gene, *rplO* (ribosomal protein L15)^22^. This approach has proven useful in several reports for examination of community structure via shotgun sequencing that often results in poor 16S ribosomal RNA gene assemblies^23–25^. Based on *rplO* genes, 466 distinct organisms were identified in the biocrust across all conditions (Supplementary Table 2). As observed for biocrust metabolites, community structure was primarily driven by time since wetting. At the phylum level, the most drastic change was a shift from a cyanobacteria-dominated community at early time points (17-28% at 3 min to 1-3% by 49.5 h) to a Firmicutes-dominated community by 49.5 h (4-5% at 3 min to 19-39% by 49.5 h) (Supplementary Figure 3). Other dominant phyla included Proteobacteria and Actinobacteria, which appeared to be indifferent to wetting (*i.e.* their relative abundance was more evenly-distributed across wetting) (Supplementary Figure 3).

In order to use previous exometabolomic studies to link soil microbe-metabolite abundances in biocrust, *rplO* gene sequences of the profiled biocrust bacterial isolates^20^ were compared to all *rplO* genes obtained from biocrust. With this approach, we identified four relatively abundant isolate-related phylotypes in the biocrust that were selected for further analyses and exometabolomics comparisons: *Microcoleus* spp. (a filamentous Cyanobacterium and primary producer), two Firmicutes (referred to here as *Anoxybacillus* sp. and *Bacillus* sp.) and *Blastococcus* sp. (an actinobacterium) (Table 1, Supplementary Figure 4). *Microcoleus* spp. is known to be a pioneer species responsible for initial soil stabilization and biocrust formation^26^ and in our study was the most dominant in early wetup biocrust, accounting for 10-25% of the entire microbial community at 3 min across all successional stages (Supplementary Figure 5). The two Firmicutes, *Anoxybacillus* sp. and *Bacillus* sp., are likely to be physically-associated with *Microcoleus* filaments^20^ and their relative abundance increased during wetting. The most abundant of these, *Anoxybacillus sp.*, was a mid-wetup responder and peaked at 9 h for successional levels A, B and D (16-24% of the community) and at 18 h for successional level C (24% of the community) (Supplementary Figure 5). *Bacillus* sp. reached its peak abundance at later time points, accounting for up to 3% of the community by 42 h in successional level C (Supplementary Figure 5), noting that by the later time points the community was less dominated by any one particular organism (Supplementary Figure 5). Finally, *Blastococcus* sp. abundance was found to be relatively resistant to wetting and was somewhat evenly distributed across all conditions (0.1-2% of the community) (Supplementary Figure 5).

**Figure 3.**
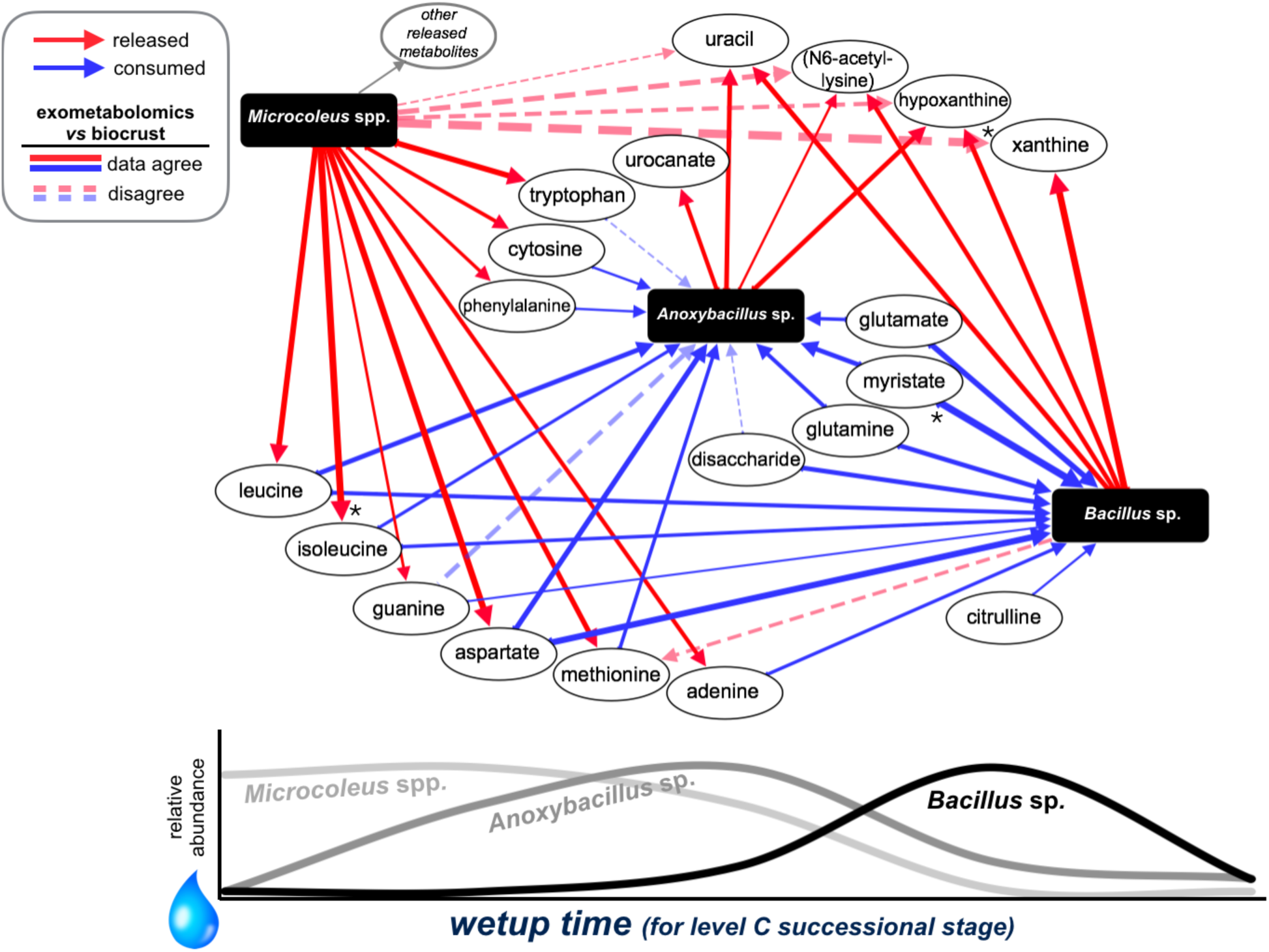
Simplified biocrust foodweb of three dominant bacteria based on exometabolomic patterns. This network displays the relationships between metabolites and three dominant organisms as they increase and decrease across wetting and successional stages in biocrust. As *Microcoleus* spp. immediately increases in relative abundance during early time points, many released metabolites (based on exometabolomics) are positively correlated with *Microcoleus* spp. (solid red arrows) and as the two relatively-abundant *Bacilli* increase (first *Anoxybacillus* sp. then *Bacillus* sp.), consumed metabolites decrease and are negatively correlated with these bacteria (solid blue arrows) and released metabolites are positively correlated (solid red arrows). Dotted arrows indicate metabolites that are released (red) or consumed (blue) that did not display the expected correlation relationship with that organism. The thickness of the line corresponds to the absolute value of the Spearman’s *rho* correlation coefficient. * FDR < 0.05 for individual microbe-metabolite correlations. The expected directionality (solid lines versus dotted lines) was significant as determined by the exact binomial test (p < 1 x 10^-5^).

### Linking microbe-metabolite abundances based on isolate exometabolomics profiling

To determine how isolate substrate preferences impacted *in situ* exometabolite composition, we evaluated microbe-metabolite correlations, focusing on metabolites known to be released or consumed by the related isolates of *Microcoleus* spp., *Anoxybacillus* sp., *Bacillus* sp. and *Blastococcus* sp. The expectation was that released metabolites would be positively correlated with the relative bacterial abundance while consumed metabolites would be negatively correlated (Figure 1B) across both wetting and successional stage. To link previous isolate exometabolomics data with the current biocrust exometabolome dataset (Figure 2), we determined the degree of correlation between the metabolites that were previously found to be consumed and released by biocrust isolates^20^ and the four relatively-abundant bacteria of interest found in the biocrust (*Microcoleus* spp., *Anoxybacillus* sp., *Bacillus* sp. and *Blastococcus* sp). Of the 85 metabolites identified in the biocrust soil water, 32 matched the isolate exometabolome dataset (Supplementary Table 3).

To assess the directionality (positive versus negative correlations) of predicted soil microbe-metabolite relationships, we performed Spearman’s rank correlation analyses. We then used an exact binomial test to evaluate the possibility that the correct directionality could occur by chance (Supplementary Table 4). Strikingly, of the 71 microbe-metabolite relationships evaluated (Supplementary Figure 6), 76% had the predicted directionality and would be very unlikely to occur by chance (two-tailed p-value < 1 x 10^-5^; Supplementary Table 4).

We next used our data to hypothesize a dynamic exometabolomic web of microbes, largely reflecting the release of metabolites by the primary producer (*Microcoleus* spp.) followed by consumption by the two heterotrophs that displayed a large degree of cycling across wetting (*Anoxybacillus* sp. and *Bacillus* sp.) (Figure 3). *Blastococcus* sp. is not shown in Figure 3 since this organism did not drastically change across time and to simplify visualization of the network. Of the large set of metabolites that were most highly-released by *M. vaginatus* PCC 9802^20^, 20 of these were detected in the biocrust soil water and most (65%) were positively correlated with *Microcoleus* spp. (Figure 3 and Supplementary Figure 6) across wetting and successional stages. While *Microcoleus* spp. was most abundant immediately following wetting, most of these metabolites (80%) reached their highest level during the first three time points (3 min, 9 h or 18 h) just after the *Microcoleus* spp. spike, suggesting release by *Microcoleus* spp. followed by increasing consumption by heterotrophs.

Consistent with a heterotrophic lifestyle, metabolites were primarily negatively correlated with the abundances of *Anoxybacillus* sp., *Bacillus* sp. and *Blastococcus* sp. Of the metabolites that were consumed by the *Anoxybacillus* sp.- related isolate, D1B51 (Table 1)^20^, 12 were detected in the current biocrust soil water samples and reached their highest level early-on (at either 3 min, 9 or 18 h), decreasing just after the peak in *Anoxybacillus* sp. Nine of these metabolites were negatively correlated with *Anoxybacillus* sp., consistent with metabolite consumption, and all four D1B51-released metabolites were positively correlated with *Anoxybacillus* sp. (Figure 3 and Supplementary Figure 6). As for the less dominant organisms, the late-wetup responder, *Bacillus* sp., was negatively correlated with all 10 metabolites that were consumed by the related isolate (L2B47) and positively correlated with 4 out of 5 isolate-released metabolites (Figure 3 and Supplementary Figure 6). Furthermore, *Blastoccocus* sp., was negatively correlated with 13 out of the 15 metabolites that were consumed by the related isolate (L1B44), while a single isolate-released metabolite was positively correlated (tryptophan; Supplementary Figure 6). Finally, the closest biocrust phylotypes of the three remaining exometabolomic-profiled isolates (L1B56, D1B2 and D1B45) accounted for 0.1% or less of the microbial community in our metagenomes and thus, not surprisingly, did not display exometabolite-based microbe-metabolite relationships (data not shown).

### Transcriptomics support the link between soil microbes and metabolites

Transcriptomics has the potential to test if gene expression is consistent with the predicted substrate utilization and release. As an initial proof of concept, we further analyzed data obtained from a previous study that evaluated *M. vaginatus* gene expression following wetup and drydown in biocrusts obtained from the same field site^27^. We found that pathways involved in the biosynthesis of amino acids (KEGG pathways ‘biosynthesis of amino acids’, ‘phenylalanine, tyrosine and tryptophan biosynthesis’ and ‘valine, leucine and isoleucine biosynthesis’) all increased dramatically during early wet-up (Supplementary Figure 6, Supplementary Table 5). In contrast, pathways involved in the degradation of these same metabolites were relatively constant (‘phenylalanine metabolism’) or only slightly increased (‘tryptophan metabolism’ and ‘valine, leucine and isoleucine biosynthesis’) following wet-up (Supplementary Figure 7) consistent with the early-increase of most amino acids in biocrust soil water in the present study and the release of these metabolites by *M. vaginatus* PCC 9802^20^.

## DISCUSSION

Sequencing has the potential to link exometabolite composition to specific microbes based on genome annotations. However, based on these data alone, relating metabolic potential to activity is challenging. Despite this, sequencing and other approaches have started to shed light on the impact individual organisms^28^, microbial genes^29^ and enzymatic activities^30^ have on the chemistry within their environment. Here, we evaluated exometabolite profiles of individual bacteria for linking soil metabolites to bacteria in biocrust, a critical ecosystem that lends itself to studies of community responses to soil wetting.

We found that a wetting event set in motion an immediate cascade of microbial activities marked by a drastic shift in community structure. The dominance of Cyanobacteria during early time points is consistent with previous reports^31^ as is the subsequent Firmicutes-bloom^9,32^. The Firmicutes phylum consists mostly of gram-positive, spore-forming bacteria with rapid generation times, enabling them to ‘bloom’ upon soil wetting^33,34^. The observed switch from a Cyanobacteria-dominated community to a Firmicutes-dominated community (mostly *Anoxybacillus* sp. in this study) agrees with our observations of metabolite release by the dominant photoautotroph (*Microcoleus* spp.)^20^ followed by consumption and growth of diverse heterotrophs (*e.g.* Firmicutes), possibly including symbiotic nitrogen-fixers^35^. While we did not observe evidence of fixed nitrogen transfer into Cyanobacteria, this process may occur during dry-down, when nitrogen-rich nutrients may be released upon the mother cell lysis stage of sporulation^36^.

It has been suggested that copiotrophic organisms (*e.g.* many Firmicutes) are superior competitors for a limited number of compounds whereas oligotrophs (*e.g.* many Actinobacteria) support a more stable population by using a wider range of substrates^37^. Our previous exometabolomics work is consistent with this view and showed that the two Firmicutes isolates depleted the narrowest range of substrates (10%) whereas the two Actinobacteria used almost twice as many^20^. Here we find that unlike the boom-bust cycle of Fimicutes, the Actinobacteria phylum (such as *Blastococcus* sp.) may be more resistant to wetting. This provides limited evidence that utilization of diverse substrates, consistent with oligotrophy, may enable slow but continuous growth under conditions with highly dynamic exometabolite pools.

The community dynamics that were caused by biocrust wetting resulted in strong microbe-metabolite relationships that were conserved from one successional stage to another for the four bacteria of interest (*Microcoleus* spp., *Anoxybacillus* sp., *Bacillus* sp. and *Blastococcus* sp.) (Supplementary Figure 8). This supports the notion that the water-soluble SOM in these biocrusts, to a large degree, originates from and is controlled by microbes^15^ and the composition of this pool may be predictable if a change in microbial community structure is anticipated. This finding has particular significance for biocrusts, since changes in temperature and rainfall are expected to shift microbial community structure^38,39^. As a result, these alterations are expected to impact SOM cycling especially if there is loss of taxa responsible for utilization or production of specific SOM components.

Next, we explored the connection between the observed microbe-metabolite relationships in biocrust and culture-based exometabolite profiles. Overall, we found that isolate exometabolomic patterns were conserved in the intact biocrust soil microbial community. The expected directionality (positive or negative microbe-metabolite correlations) (solid arrows in Figure 3) was significantly higher than predicted by chance, indicating a linkage between laboratory observations and *in situ* soil activities. While most metabolites displayed the expected patterns, some biocrust soil water metabolites (*i.e.* uracil, N6-acetyl-lysine, hypoxanthine and xanthine) were inconsistent with *M. vaginatus* PCC 9802 exometabolite profiles. However, these were also released by and positively correlated with *Bacillus* sp. Deconvoluting this may be possible using dynamic utilization models^40,41^ to account for the relative contributions of the two organisms. Ultimately, this same approach could be used to account for rare community members that may also have an impact on the exometabolite pool or may alter the metabolism of other microbes^42,43^. Although outside of the scope of the current work, we anticipate that these substrate-genome linkages could be further tested and refined by using other approaches. Stable isotope probing coupled with labeled DNA sequencing^35,44^ and integrated NanoSIMS and FISH imaging^45,46^ may be used to examine the spatial localization of microbes and their activities.

We next used the biocrust microbe-metabolite relationships to postulate a dynamic exometabolomics web describing the wetting response of three dominant bacteria in the biocrust (Figure 3). This network displays the release of many metabolites, especially amino acids, by *Microcoleus* spp. followed by consumption by the two heterotrophic Firmicutes. This suggests unique organismal roles in the biocrust foodweb including the preferential consumption of aromatic amino acids (tryptophan and phenylalanine) by *Anoxybacillus* sp. and branched-chain amino acids (leucine and isoleucine) by both Firmicutes. Interestingly, we also observe that these Firmicutes release nucleobases (uracil, hypoxanthine and xanthine), consistent with our earlier reports of heterotrophs releasing these compounds^47^. This may reflect a nitrogen-scavenging mechanism by consuming N-containing substrates (cytosine, adenine, guanine and histidine), producing uracil, hypoxanthine, xanthine and urocanate as byproducts. Knowledge of these functional linkages between metabolites and microbes has the potential to help understand and predict nutrient cycling in terrestrial microbial ecosystems^48^ analogously to the many organisms that have been linked to specific transformations within marine ecosystems. For example Cyanobacteria release and reuptake organic carbon^49^, a variety of uncultured taxa utilize dissolved proteins^44^ and SAR11 bacteria assimilate amino acids and dimethylsulfoniopropionate^50^.

We attribute much of the success of this study to the suitability of the biocrust ecosystem. One such advantage is that biocrust soil in this study is primarily quartz sand, facilitating metabolite analysis compared to many other soils which are typically rich in clays and other strongly-sorptive mineral surfaces^51^. Accurately representing the competition between microbes and mineral surfaces would require additional studies examining mineral-metabolite sorption dynamics^52,53^. Another relatively simplifying factor is that the biocrust community, unlike many other soils, is dominated by a few bacteria, greatly enabling accurate correlations between taxa and metabolites. Furthermore, there is a general lack of consensus of isolate-to-community comparisons and what constitutes a valid comparison especially with the use of ribosomal protein genes as phylogenetic markers. Exometabolite-profiled isolates and their related biocrust phylotypes ranged between 86.3-92.0% identical in their *rplO* sequence (Table 1), likely placing them in the same genus, and with more certainty, the same family. Despite the low taxonomic resolution, the observed functional similarity, agrees with reports suggesting that metabolic traits are largely conserved at the phylum level^54^. We anticipate that in order to accurately predict microbe-metabolite relationships for more diverse communities and complex environments, a large number of relevant taxa would need to be subjected to exometabolite profiling. Accounting for switching between metabolic states will require profiling under diverse environmental conditions. For example, the discrepancy between metabolites that were released by *M. vaginatus* PCC9802^20^, but were not correlated with *Microcoleus* spp. abundance in the present study may be due to different metabolic processes occurring during the day (photosynthesis) versus night (respiration) (Diel cycle)^27,55^. Thus, modeling approaches will be required to account for metabolic state switching among other processes. One exciting possibility of expanded exometabolomic datasets, is that knowledge of uptake and release of metabolites can be used as boundary constraints for flux-balance analysis^56^ and trait-based models^57^ providing a genome-scale approach for linking soil metabolites with metagenomic data. For example, OptCom^58^, a multi-level and multi-objective flux balance analysis framework to understand metabolism within microbial communities, which currently primarily relies on genomic information, could be used in conjunction with exometabolomic data.

In conclusion, this study shows that isolate exometabolite patterns are conserved within an intact biocrust community, relating community structure and metabolite composition. We expect that exometabolomic characterization of additional taxa and determination of mineral-metabolite sorption dynamics, under a range of environmentally relevant conditions (e.g. day/night cycles), integrated with modeling approaches will further enhance the predictive power of these relationships. These studies may help pave the way for interpretation and use of metagenomic and metatranscriptomic approaches for linking soil chemistry to soil microbiomes to define exometabolite webs of microbes in complex ecosystems.

## MATERIALS AND METHODS

### Materials

LC/MS-grade water and LC/MS-grade methanol (CAS 67-56-1) were from Honeywell Burdick & Jackson (Morristown, NJ). LC/MS-grade acetonitrile (CAS 75-05-8) and ammonium acetate (CAS 631-61-8) were from Sigma-Aldrich (St. Louis, MO). LC/MS internal standards included MOPS (CAS 1132-61-2), HEPES (CAS 7365-45-9), 3,6-dihydroxy-4-methylpyridazine (CAS 5754-18-7), 4-(3,3-dimethyl-ureido)benzoic acid (CAS 91880-51-2), d_5_-benzoic acid (CAS 1079-02-3) and 9-anthracenecarboxylic acid (CAS 723-62-6) from Sigma-Aldrich.

### Sample collection

Petri dishes (6 cm^2^ x 1 cm depth) were used to core biocrust samples from the Green Butte Site near Canyonlands National Park (38°42'54.1"N, 109°41'27.0"W, Moab, UT, USA). Samples were collected along an apparent maturity gradient of Cyanobacteria-dominated biocrusts ranging from light, young (level A) to darker, more mature (level D) (Supplementary Figure 1). Samples were air-dried in the field and brought back to the laboratory where they were maintained in a dark desiccation chamber.

### Biocrust wetting

Biocrust (0.5 g) was transferred to each well within 12-well plates. Sterile LC/MS-grade water (1 mL) was added to each sample and placed under a 12 h light (~300 µmol photons/m^2^s)/ 12 h dark cycle. Microcosms were completely enclosed by aluminum foil to prevent infiltration by outside light sources. At each time point (3 min, 9 h, 18 h, 42 h and 49.5 h), biocrust and soil water were removed and placed in 2 mL Eppendorf tubes and 500 µL of additional water was used to rinse out the wells and added to the sample. Tubes were centrifuged at 5000 x *g* for 5 min and supernatant (biocrust soil water) was pipetted and placed in new 2 mL tubes. Remaining biocrust was stored at −80°C until nucleic acid extraction was performed. There were five replicates, five time points and four successional stages of crust resulting in 100 total samples.

### Metabolite extraction and LC/MS analysis

Biocrust soil water samples (1.5 mL) were lyophilized and resuspended in methanol (200 µL) containing internal standards (2-10 µg/mL) and filtered through 96-well Millipore filter plates (0.2 µm PVDF) by centrifuging at 1500 x *g* for 2 min. Samples were analyzed using normal-phase LC/MS with a ZIC-pHILIC column (150 mm × 2.1 mm, 3.5 µm 200 Å, Merck Sequant, Darmstadt, Germany) using an Agilent 1290 series UHPLC (Agilent Technologies, Santa Clara, California, USA). Chromatographic separation was achieved using two mobile phases, 5 mM ammonium acetate in water (A) and 90% acetonitrile w/ 5 mM ammonium acetate (B) at a flow rate of 0.25 mL/min with the following gradient: 100% B for 1.5 min, linear decrease to 50% B by 25 min, held until 29.9 min then returned to initial conditions by 30 min with a total runtime of 40 min. Column temperature was maintained at 40°C. For MS, negative mode data were acquired on an Agilent 6550 quadrupole time-of-flight mass spectrometer and positive mode data were acquired on a Thermo QExactive (Thermo Fisher Scientific, Waltham, MA). Fragmentation spectra (MS/MS) were acquired for some metabolites using collision energies of 10-40 eV.

Metabolomics data were analyzed using Metabolite Atlas^59^ in conjunction with the Python programming language. Internal standards were assessed from each sample to ensure peak area and retention times were consistent from sample-to-sample. Quality control mixtures were included at the beginning, end and throughout the runs to ensure proper instrument performance (*m/z* accuracy and retention time and peak area stability). Sample QC failed for some replicates including all 9 h wetup level D samples and were not included for further analyses. Metabolite identifications were based on two orthogonal data relative to authentic standards and/or the Metlin database^60,61^ and are provided in Supplementary Table 1. Putative identifications were assigned in the cases where these criteria were not met and are indicated by parentheses in figures. To explore the degree of variation in biocrust metabolite profiles across wetting and successional stages, biocrust samples were PCA-ordinated based on their metabolite profiles.

### DNA extraction, sequencing and microbial annotation

DNA was extracted from biocrust (0.25 g) using the MoBio Powersoil DNA isolation kit (MoBio Laboratories, Inc, Carlsbad, CA) resulting in 100 uL of eluted DNA. Library preparation and sequencing were done at the QB3 facility at the University of California, Berkeley using Illumina HiSeq4000 (see supplementary methods for details on metagenome analysis). In recent studies^23-25^, ribosomal protein genes have been used as phylogenetic markers as an alternative to the more classical 16S ribosomal RNA gene. Ribosomal protein genes exist as single copies in almost all genomes, assemble well from metagenome datasets, are well-conserved and have produced higher resolution phylogenetic trees^23^. Given these advantages, the 50S ribosomal protein L15 (*rplO*) gene was well-represented in our metagenomes and was therefore used as a phylogenetic marker to examine the relative abundance of individual organisms within the microbial community across wetting and successional stages. The *rplO* genes from the genomes of the seven exometabolite profiled biocrust bacterial isolates^20^ were compared to all the *rplO* genes recovered from biocrust. Those with the highest percent similarity were considered the “closest relatives” to the isolates and are reported in Table 1.

### Microbe-metabolite correlations and statistics

Correlations were used to identify microbe-metabolite relationships across both wetting and successional stages. Spearman’s rank (*rho*) correlation coefficients for every pairwise (microbe-metabolite) relationship and p-values (unadjusted and FDR-adjusted) were calculated using the cor() stats function in R. A Spearman’s *rho* value greater than or equal to 0.5 was considered “highly correlated” and less than or equal to −0.5 was considered “highly negatively correlated”. To test if the overall observed directionality (positive versus negative correlations) was due to chance rather than as would be predicted based on exometabolomics (release versus consumption), the exact binomial test was conducted using R (binom.test) with a total of 71 “trials” or observed microbe-metabolite interactions (Supplemental table 4).

### Exometabolomics comparison and analysis

Of the 85 metabolites detected in biocrust soil water, 32 were previously analyzed for consumption and release by biocrust isolates^20^. For continued analyses of those data here, fold-change was calculated by dividing the average peak area of each metabolite in (isolate) inoculated spent media by the non-inoculated control spent media (raw data can be found in the Supplemental Table in Baran et al, 2015). A metabolite was considered consumed if the fold-change was 0.5 or less and released metabolites had a fold-change of 2 or greater.

### Microcoleus gene expression analysis

*Microcoleus* genes from Supplementary Table S6 from Rajeev et al (2013) were categorized into KEGG pathways (for genes containing KEGG ID numbers). Analyses focused on pathways that are primarily anabolic or catabolic for metabolites that were released by *Microcoleus* PCC 9802. The average fold-change (relative to dry biocrusts) and standard errors were calculated for all genes belonging to pathways of interest.

## ACKNOWLEDGEMENTS

This work was funded by the Office of Science Early Career Research Program, Office of Biological and Environmental Research, of the U. S. Department of Energy under contract number DE-AC02-05CH11231. DNA was sequenced using the Vincent J. Coates Genomics Sequencing Laboratory at UC Berkeley, supported by NIH S10 OD018174 Instrumentation Grant. We thank Rebecca Lau for technical assistance in biocrust sample collection and experimentation.

## AUTHOR CONTRIBUTIONS

T. L. S. and T. R. N. conceived the study, designed the experiments and wrote the manuscript. T. L. S. performed the experiments. T. L. S. and B. P. B. analyzed the metabolomics data. U. K. analyzed the metagenomics data. T. L. S. and J. M. S. conducted correlation and statistical analyses. All co-authors commented on the design of experiments, data analysis and draft manuscripts.

## REFERENCES

1. Paustian, K. et al. Climate-smart soils. Nature 532, 49–57 (2016).

2. Friedlingstein, P. et al. Climate–carbon cycle feedback analysis: Results from the C4MIP model intercomparison. J. Climate 19, 3337–3353 (2006).

3. Judd, K. E., Crump, B. C. & Kling, G. W. Variation in dissolved organic matter controls bacterial production and community composition. Ecology 87, 2068–2079 (2006).

4. Collins, S. L. et al. A multiscale, hierarchical model of pulse dynamics in arid-land ecosystems. Annu. Rev. Ecol. Evol. Syst. 45, 397–419 (2014).

5. Siciliano, S. D. et al. Soil fertility is associated with fungal and bacterial richness, whereas pH is associated with community composition in polar soil microbial communities. Soil Biol. Biochem. 78, 10–20 (2014).

6. Berthrong, S. T., Buckley, D. H. & Drinkwater, L. E. Agricultural management and labile carbon additions affect soil microbial community structure and interact with carbon and nitrogen cycling. Microb. Ecol. 66, 158–170 (2013).

7. Maestre, F. T. et al. Increasing aridity reduces soil microbial diversity and abundance in global drylands. Proc Natl Acad Sci USA 112, 15684–15689 (2015).

8. Barnard, R. L., Osborne, C. A. & Firestone, M. K. Responses of soil bacterial and fungal communities to extreme desiccation and rewetting. ISME J 7, 2229–2241 (2013).

9. Placella, S. A., Brodie, E. L. & Firestone, M. K. Rainfall-induced carbon dioxide pulses result from sequential resuscitation of phylogenetically clustered microbial groups. Proc Natl Acad Sci USA 109, 10931–10936 (2012).

10. Belnap, J., Weber, B. & Büdel, B. Biological Soil Crusts as an Organizing Principle in Drylands. in Biological Soil Crusts: An Organizing Principle in Drylands (eds. Weber, B., Büdel, B., & Belnap, J.,) 3–13 (Springer, 2016).

11. Maestre, F. T. et al. Structure and functioning of dryland ecosystems in a changing world. Annu. Rev. Ecol. Evol. Syst. 47, 215–237 (2016).

12. Elbert, W. et al. Contribution of cryptogamic covers to the global cycles of carbon and nitrogen. Nat. Geosci. 5, 459–462 (2012).

13. Garcia-Pichel, F. & Pringault, O. Cyanobacteria track water in desert soils. Nature 413, 380–381 (2001).

14. Baldock, J. A. & Nelson, P. N. Soil Organic Matter. in Handbook of Soil Science (ed. Sumner, M. E.) B25–B84 (CRC Press, 2000).

15. Schmidt, M. W. I. et al. Persistence of soil organic matter as an ecosystem property. Nature 478, 49–56 (2011).

16. Kogel-Knabner, I. The macromolecular organic composition of plant and microbial residues as inputs to soil organic matter. Soil Biol. Biochem. 34, 139–162 (2002).

17. Swenson, T. L., Jenkins, S., Bowen, B. P. & Northen, T. R. Untargeted soil metabolomics methods for analysis of extractable organic matter. Soil Biol. Biochem. 80, 189–198 (2015).

18. Warren, C. R. Comparison of methods for extraction of organic N monomers from soil microbial biomass. Soil Biol. Biochem. 81, 67–76 (2015).

19. Fierer, N. & Lennon, J. T. The generation and maintenance of diversity in microbial communities. Am. J. Bot. 98, 439–448 (2011).

20. Baran, R. et al. Exometabolite niche partitioning among sympatric soil bacteria. Nat. Commun. 6, 1–9 (2015).

21. Silva, L. P. & Northen, T. R. Exometabolomics and MSI: deconstructing how cells interact to transform their small molecule environment. Curr. Opin. Biotechnol. 34, 209–216 (2015).

22. Yutin, N., Puigbo, P., Koonin, E. V. & Wolf, Y. I. Phylogenomics of prokaryotic ribosomal proteins. PLoS ONE 7, (2012).

23. Hug, L. A. et al. A new view of the tree of life. Nat. Microbiol. 1, 1–6 (2016).

24. Wu, M. & Eisen, J. A. A simple, fast, and accurate method of phylogenomic inference. Genome Biol. 9, R151 (2008).

25. Sharon, I. et al. Accurate, multi-kb reads resolve complex populations and detect rare microorganisms. Genome Res. 25, 534–543 (2015).

26. Garcia-Pichel, F. & Wojciechowski, M. F. The Evolution of a Capacity to Build Supra-Cellular Ropes Enabled Filamentous Cyanobacteria to Colonize Highly Erodible Substrates. PLoS ONE 4, e7801–6 (2009).

27. Rajeev, L. et al. Dynamic cyanobacterial response to hydration and dehydration in a desert biological soil crust. ISME J 7, 2178–2191 (2013).

28. Walsh, A. M. et al. Microbial succession and flavor production in the fermented dairy beverage kefir. mSystems 1, e00052–16 (2016).

29. Ding, J. et al. Integrated metagenomics and network analysis of soil microbial community of the forest timberline. Sci. Rep. 5, 7994–10 (2015).

30. Trivedi, P. et al. Microbial regulation of the soil carbon cycle: Evidence from gene-enzyme relationships. ISME J 10, 2593–2604 (2016).

31. Garcia-Pichel, F., Lopez-Cortes, A. & Nubel, U. Phylogenetic and morphological diversity of cyanobacteria in soil desert crusts from the Colorado Plateau. Appl. Environ. Microbiol. 67, 1902–1910 (2001).

32. Angel, R. & Conrad, R. Elucidating the microbial resuscitation cascade in biological soil crusts following a simulated rain event. Environ. Microbiol. 15, 2799–2815 (2013).

33. Nicholson, W. L., Munakata, N., Horneck, G., Melosh, H. J. & Setlow, P. Resistance of Bacillus endospores to extreme terrestrial and extraterrestrial environments. Microbiol Mol Biol Rev 64, 548–572 (2000).

34. Nicholson, W. L. Roles of Bacillus endospores in the environment. Cell. Mol. Life Sci. 59, 410–416 (2002).

35. Pepe-Ranney, C. et al. Non-cyanobacterial diazotrophs mediate dinitrogen fixation in biological soil crusts during early crust formation. ISME J 10, 287–298 (2016).

36. Hosoya, S., Lu, Z., Ozaki, Y., Takeuchi, M. & Sato, T. Cytological analysis of the mother cell death process during sporulation in bacillus subtilis. J. Bacteriol. 189, 2561–2565 (2007).

37. Upton, A. C. & Nedwell, D. B. Nutritional flexibility of oligotrophic and copiotrophic antarctic bacteria with respect to organic substrates. FEMS Microbiol Ecol 62, 1–6 (1989).

38. Garcia-Pichel, F., Loza, V., Marusenko, Y., Mateo, P. & Potrafka, R. M. Temperature drives the continental-scale distribution of key microbes in topsoil communities. Science 340, 1574–1577 (2013).

39. Ferrenberg, S., Reed, S. C. & Belnap, J. Climate change and physical disturbance cause similar community shifts in biological soil crusts. Proc Natl Acad Sci USA 112, 12116–12121 (2015).

40. Erbilgin, O. et al. Dynamic substrate preferences and predicted metabolic properties of a simple microbial consortium. BMC Bioinformatics accepted (2017).

41. Behrends, V., Ebbels, T. M. D., Williams, H. D. & Bundy, J. G. Time-resolved metabolic footprinting for nonlinear modeling of bacterial substrate utilization. Appl. Environ. Microbiol. 75, 2453–2463 (2009).

42. Seth, E. C. & Taga, M. E. Nutrient cross-feeding in the microbial world. Front Microbiol 5, 1–6 (2014).

43. Ponomarova, O. & Patil, K. R. Metabolic interactions in microbial communities: untangling the Gordian knot. Curr. Opin. Microbiol. 27, 37–44 (2015).

44. Orsi, W. D. et al. Diverse, uncultivated bacteria and archaea underlying the cycling of dissolved protein in the ocean. ISME J 10, 2158–2173 (2016).

45. Fike, D. A., Gammon, C. L., Ziebis, W. & Orphan, V. J. Micron-scale mapping of sulfur cycling across the oxycline of a cyanobacterial mat: a paired nanoSIMS and CARD-FISH approach. ISME J 2, 749–759 (2008).

46. Woebken, D. et al. Revisiting N2 fixation in Guerrero Negro intertidal microbial mats with a functional single-cell approach. ISME J 9, 485–496 (2015).

47. Baran, R. et al. Metabolic footprinting of mutant libraries to map metabolite utilization to genotype. ACS Chem. Biol. 8, 189–199 (2013).

48. Gougoulias, C., Clark, J. M. & Shaw, L. J. The role of soil microbes in the global carbon cycle: tracking the below-ground microbial processing of plant-derived carbon for manipulating carbon dynamics in agricultural systems. J. Sci. Food Agric. 94, 2362–2371 (2014).

49. Stuart, R. K. et al. Cyanobacterial reuse of extracellular organic carbon in microbial mats. ISME J 10, 1240–1251 (2016).

50. Malmstrom, R. R., Kiene, R. P., Cottrell, M. T. & Kirchman, D. L. Contribution of SAR11 bacteria to dissolved dimethylsulfoniopropionate and amino acid uptake in the North Atlantic Ocean. Appl. Environ. Microbiol. 70, 4129–4135 (2004).

51. Kaiser, M. et al. The influence of mineral characteristics on organic matter content, composition, and stability of topsoils under long-term arable and forest land use. J. Geophys. Res. 117, G02018 (2012).

52. Swenson, T. L., Bowen, B. P., Nico, P. S. & Northen, T. R. Competitive sorption of microbial metabolites on an iron oxide mineral. Soil Biol. Biochem. 90, 34–41 (2015).

53. Lv, J. et al. Molecular-scale investigation with ESI-FT-ICR-MS on fractionation of dissolved organic matter induced by adsorption on iron oxyhydroxides. Environ. Sci. Technol. 50, 2328–2336 (2016).

54. Martiny, J. B. H., Jones, S. E., Lennon, J. T. & Martiny, A. C. Microbiomes in light of traits: A phylogenetic perspective. Science 350, aac9323–1 (2015).

55. Diamond, S., Jun, D., Rubin, B. E. & Golden, S. S. The circadian oscillator in Synechococcus elongatus controls metabolite partitioning during diurnal growth. Proc Natl Acad Sci USA 112, E1916–E1925 (2015).

56. Orth, J. D., Thiele, I. & Palsson, B. Ø. What is flux balance analysis? Nat. Biotechnol. 28, 245–248 (2010).

57. Allison, S. D. A trait-based approach for modelling microbial litter decomposition. Ecol. Lett. 15, 1058–1070 (2012).

58. Zomorrodi, A. R. & Maranas, C. D. OptCom: A multi-level optimization framework for the metabolic modeling and analysis of microbial communities. PLoS Comput. Biol. 8, e1002363 (2012).

59. Yao, Y. et al. Analysis of Metabolomics Datasets with High-Performance Computing and Metabolite Atlases. Metabolites 5, 431–442 (2015).

60. Smith, C. A. et al. METLIN: A metabolite mass spectral database. Ther. Drug Monit. 27, 747–751 (2005).

61. Sumner, L. W. et al. Proposed minimum reporting standards for chemical analysis. Metabolomics 3, 211–221 (2007).

